# An assessment of the evidence for antibacterial activity of stinging nettle (*Urtica dioica*) extracts

**DOI:** 10.1101/2021.11.17.468953

**Authors:** Freya Harrison, Jessica Furner-Pardoe, Erin Connelly

**Affiliations:** School of Life Sciences, Gibbet Hill Campus, University of Warwick, Coventry CV4 7AL, UK; Warwick Medical School, Gibbet Hill Campus, University of Warwick, Coventry CV4 7AL, UK

**Author notes:** Corresponding authors: FH & EC.

## Abstract

Stinging nettles (*Urtica* spp.) have been used in a diverse range of traditional and historical medicines from around the world for the treatment of skin diseases, wounds, urinary disorders, respiratory diseases, bone and joint pain, anaemia and other circulatory problems, as well as in cosmetic preparations for skin and haircare. As part of an interdisciplinary exploration of nettle-based remedies, we performed a systematic review of published evidence for the antimicrobial activity of *Urtica* spp. extracts against bacteria and fungi that commonly cause skin, soft tissue and respiratory infections. We focussed on studies in which minimum inhibitory concentration (MIC) assays of *U. dioica* were conducted on the common bacterial opportunistic pathogens *Escherichia coli*, *Pseudomonas aeruginosa*, *Klebsiella pneumoniae* and *Staphylococcus aureus*. No studies used fresh leaves (all were dried prior to use), and no studies prepared nettles in weak acid (corresponding to vinegar) or in fats/oils, which are common combinations in historical and traditional preparations. We addressed this gap by conducting new antibacterial tests of extracts of fresh *U. dioica* leaves prepared in vinegar, butter or olive oil against *P. aeruginosa* and *S. aureus*. Our systematic review and additional experimental data leads us to conclude that there is no strong evidence for nettles containing molecules with clinically useful antimicrobial activity. It seems most likely that the utility of nettles in traditional topical preparations for wounds may simply be as a “safe” absorbent medium for keeping antibacterial (vinegar) or emollient (oils) ingredients at the treatment site.

**Data Summary:** All data relating to the systematic review are supplied in the Datasets and Supplementary tables contained in the Supporting Data file.

## Introduction

Stinging nettles (*Urtica* species including *U. dioica*, *U. urens*, *U. pilulifera*) have a broad geographical distribution, and have been used in a range of traditional and historical medicines from around the world. Various parts of nettle plants appear in preparations for the treatment of skin diseases, urinary disorders, respiratory diseases, bone and joint pain, anaemia and other circulatory problems, as well as in cosmetic preparations for skin and haircare, in traditional remedies or historical medical texts from many cultures (1–4). Some of these uses appear rational given current scientific knowledge. For instance, nettles are effective accumulators of heavy metals and are generally agreed to be a good source of iron, so their use in anaemia treatments appears sensible (5). Additionally, the anti-inflammatory potential of natural products from nettles has received significant research attention, and some studies have suggested evidence that nettles could provide new treatments for rheumatoid arthritis and osteoarthritis (1, 6, 7). Nettles also have a long history of safe incorporation into food and drink (e.g. nettle stews, cheeses, beers) and topical cosmetic products such as shampoo and soap.

Given the widespread historical medical and culinary use of nettles, as well as their long growing season and lack of conservation concern, it is unsurprising that many researchers have also assessed the antimicrobial properties of nettles. Recent narrative reviews bring together a wide body of experimental work assessing the antibacterial or antifungal activity of nettle extracts (1, 6, 7). However, there has been no systematic review or meta-analysis of published work on the antimicrobial properties of nettles and the primary research does not appear to have generated any useful clinical leads. As of August 2021, The Cochrane Central Register of Controlled Trials included 111 clinical trials containing “urtica” in their keyword, title or abstract. These included trials of nettle sting, nettle extracts or compound natural product preparations/nutraceuticals for conditions including acne, burns, wound healing, joint pain, arthritis, urinary symptoms of benign prostatic hyperplasia, menstrual or post-partum bleeding, diabetes, allergic rhinitis and even COVID-19. Trials of nettle products for antibacterial or antifungal activity were conspicuous by their almost total absence.

As part of an interdisciplinary study of European historical medicine, we created a dataset of as many nettle-based remedies as possible from select early and late medieval sources for comparison of ingredients and recipe structure across a wide span of time and texts. The dataset, along with full project details, will shortly be published (8) and also made available at http://ancientbiotics.co.uk. The species include the common stinging nettle (*U. dioica*) and other stinging *Urtica* species. The medieval texts examined here also mention dead-nettles (white or red), which may be translated as *Lamium* spp. In these medieval texts, *nettle* can be any stinging plant of *Urtica* spp. or *Lamium* spp., but the term is especially connected with *U. dioica*. The final dataset contains 139 recipes from 30 sources (Old English, Middle English, Latin, and medieval Welsh) from the ninth to the fifteenth century.

Types of remedies include plasters, suppositories, gargles, drinks, salves, ointments, and external cleansing washes for many disease conditions such as dog bites, inflammation, post-surgical swellings, pustules, restraining blood (nosebleeds, wounds, menstrual, surgical wounds), healing and cleaning rotten wounds, swellings and abscesses of the mouth and throat, aches and pains, gout, dropsy, fevers, colic, cough, respiratory disorders, urinary disorders, “leprosy” and skin disorders, bruises and broken bones. Nettles remedies typically specify the part of the plant being used, such as juice, seeds, root, powder, leaves, flowers or the whole plant. This long list of preparations and conditions appears to suggest that nettles are being used as a generic cure-all for all kinds of internal and external issues; however, in this particular dataset, nettles are concentrated in applications for wound care in combination with vinegar, wine, salt, or fats and oils. In both early and late medieval recipes, often nettles are pounded together with just one of those ingredients and applied to the wound. In other cases, nettles appear in combination with other medicinal ingredients such as this apparently general-purpose ointment from an early medieval text:

> A wound salve, take sprigs of woad and nettle as well, pound well, boil in butter and strain through a cloth, add white salt and whisk well. (9)

And the following example from a later medieval hunting text for the treatment of “men and beasts bitten by mad hounds”:

> For men may make a sauce of salt, vinegar, and strong garlic, peeled and crushed, and nettles together, and also hot as it may be suffered, to lay upon the bite. And this is a good medicine and a true, for it has been proved. And every day it should be laid upon the bite two times, also hot as it may be suffered, until it is healed or else by nine days. And yet there is another medicine better than all the others. Take leeks, and strong garlic, and spring onion, and rue, and nettles, and cut them small with a knife. And then mix them with olive oil and vinegar and boil them together. And then take all the herbs, also hot as they may be suffered, and lay upon the wound every day two times until the wound be well healed or at the least by nine days. (10)

Vinegar contains acetic acid, which is known to have antimicrobial activity at low concentrations (11, 12), and a clinical trial is currently in progress in the United Kingdom to assess optimal dosing of acetic acid as a topical treatment for burn wound infections (13). The antimicrobial activity of fats (butter, lard, etc.) and oil is not known nor do these ingredients immediately suggest antimicrobial efficacy. Fats and oils may be functioning primarily as palliative and binding agents (perhaps even blood restraint) for healing ointments in wound care.

In our humanities-led exploration of nettle-based remedies, we aimed to identify any patterns in medical practice (the parts of plant used, the preparation methods, combination with other ingredients). In the present article, we aim to place this research in the context of published studies on the activity of nettle extracts on common wound pathogens by undertaking a systematic review of published evidence for the antimicrobial activity of *Urtica* spp. extracts against bacteria and fungi that commonly cause skin, soft tissue and respiratory infections. A variety of *Urtica* species, pathogen species, preparation methods and assay methods were used. To be able to compare activity across studies and preparation methods, we focussed on studies in which minimum inhibitory concentration (MIC) assays of *U. dioica* were conducted on the common bacterial opportunistic pathogens *Escherichia coli*, *Pseudomonas aeruginosa*, *Klebsiella pneumoniae* and *Staphylococcus aureus*. The published studies were of variable quality, and we did not find strong evidence for antibacterial activity. No studies used fresh leaves (all were dried prior to use), and no studies prepared nettles in weak acid (corresponding to vinegar) or in fats/oils. We addressed this gap by conducting new antibacterial tests of extracts of fresh *U. dioica* leaves prepared in vinegar, butter or olive oil against *P. aeruginosa* and *S. aureus* and again found no evidence of antibacterial activity.

Our systematic review and additional experimental data leads us to conclude that there is no strong evidence for nettles containing molecules with clinically useful antimicrobial activity. It is possible that *U. dioica* does in fact contain effective antimicrobial natural products, but they have not yet been reproducibly extracted in laboratory conditions. Future studies may yet identify interesting antimicrobial compounds from nettles, and reproducible methods for extracting them. We have not addressed any immunomodulatory potential of nettle-derived natural products in the present study, but there are published data which suggest that nettle extracts could possess anti-inflammatory activity. However, it seems most likely that the utility of nettles in traditional topical preparations for wounds may simply be as a “safe” absorbent medium for keeping antibacterial (vinegar) or emollient (oils) ingredients at the treatment site.

## Methods

### Systematic review

Searches for primary literature on the antimicrobial activity of *Urtica* spp. were performed in December 2020 and repeated in July 2021 using Scopus.com, PubMed, Web of Science Core Collection and the Cochrane Central Register of Controlled Trials. For each database, we searched for the Boolean string ((nettle* OR urtica) + (antimicrobial OR anti-microbial OR antibacterial OR anti-bacterial OR antifungal OR anti-fungal OR antiparasit* OR anti-parasit*)) in the document title, abstract or keywords (n.b. PubMed does not include the option to search keywords). We included documents written in any language and did not restrict by publication date. Results were then restricted to exclude reviews, editorials and commentary articles. The outputs from each search were combined in a spreadsheet. We also extracted all primary research articles included in a recent narrative review of nettle antimicrobial activity by Kregiel et al. (1) and added these to the spreadsheet. Dereplication was performed manually in Microsoft Excel using the sort function on article titles, DOIs and author names. Using three sorting procedures was important because minor differences in formatting the text of the title in the different databases meant that some duplicates were missed on the first sort, and because not all records had a DOI recorded an alternative second sorting mechanism was required for these.

Abstracts were then screened for relevance. If the abstract indicated that the full text did not contain quantitative, empirical data on the antimicrobial activity of an *Urtica* species against a known animal pathogen or *Bacillus subtilis* (as this species is commonly used as a model Gram-positive bacterium for drug testing), the record was excluded and exclusion reasons recorded. We excluded studies where nettles were only tested in combination with another substance (as we would not be able to extract data on the effect of nettle alone); where a single purified compound derived from nettles was studied; studies of nettle honey; studies of fermented nettles; studies of nettle-associate microbes for antimicrobial activity; and studies where nettles were used in green synthesis procedures. Studies were also excluded if no abstract was available. EC and FH each screened 50% of abstracts (assigned at random); 10% of abstracts were selected at random and re-screened by the other author to ensure consistent rules were applied to inclusion/exclusion.

Full text articles were sought for each of the records included at the abstract screening stage. We then excluded records for which no full text could be sourced (we emailed all corresponding authors of records for which we did not have full text access through our institutions or through open-access publication, to request articles). Full text articles were then read and assessed for eligibility. Eligible articles were those that contained quantitative data on the antimicrobial activity of nettle extracts, in the absence of any other test substance, on animal-pathogenic bacteria or *B. subtilis* expressed as a minimum inhibitory concentration (MIC) or a zone of inhibition (ZOI) diameter. Additional primary research articles referenced in these studies were added to the database and similarly screened. EC and FH collaboratively screened two full-text articles to ensure comparable processes were used, and then each screened 50% of the remaining full-text articles (assigned at random). Dr Sheyda Azimi kindly read and extracted data from two articles written in Farsi, and Prof. Orkun Soyer kindly read two articles written in Turkish and determined that we could not extract data from them, as none of the authors speaks either of these languages.

Data was extracted from all articles included at this stage. Each test of a specific concentration of a specific nettle extract on a bacterial strain was recorded in one row of a spreadsheet. Columns contained information on the article the data came from, the *Urtica* species used; the part of the plant used to make the extract; whether this material was used fresh/dried/freeze-dried; brief details on the type of extract (e.g. “crude extract – water”, “partitioned extract – chloroform”, “bioassay guided subfraction from hexane extract” etc.); brief text details of the extraction/preparation method; the temperature at which the preparation was made; the extraction time; whether the test microbe was a bacterium or a fungus (no studies were performed on eukaryotic parasites); the Gram type for bacteria; the microbial species name; details of the strain used; the assay method(s) used; and whether any standard diagnostic guidelines were followed to conduct the assay. If MIC data were reported, the units, stated value, positive and negative controls were recorded. If ZOI data were reported, the units, stated value, variance or standard deviation, number of replicates used, and positive and negative controls were recorded, along with a note on whether the reported ZOI was reported to be significantly different from any negative control in the original study. The location of recorded data in the original publication (Figure or Table number(s)) was then recorded. This spreadsheet of studies was used in further analysis.

### Plant material used in laboratory experiments

For each experiment presented below, a mix of old (lower down the stem) and young (tips of stem) leaves from *U. dioica* plants was picked in August-September from the University of Warwick’s Gibbet Hill Campus in Coventry, UK. Although it was late in the growing season, the smallest leaves at the tips of the plants were very young, being < 1 cm^2^ in area. (We note that a fifteenth-century Middle English translation of *De viribus herbarum* states that August is the best time to gather abundant nettles for medicinal purposes (14, 15)).

### Preparation of fresh nettle leaf extracts for MIC/MBC testing

*U. dioica* leaves were immediately transferred to the laboratory and chopped roughly using a sterilised knife and chopping board. The leaves were placed in Carlton Germicidal Cabinet and exposed to shortwave UV light for 10 minutes to reduce surface microbial contamination (leaves were agitated halfway through the exposure time to maximise exposure to UV by turning them over and reducing overlap between leaf pieces). 2.5 g aliquots of chopped leaves were placed into 6 sterile 50 ml screw-cap tubes, each containing 10 ¼” ceramic beads (MP Biomedicals). 5 ml red wine vinegar (Waitrose Essentials), 5 ml olive oil (Olive Branch) and 5 g unsalted British butter (Marks & Spencer) were added to two tubes each. Tubes were vortexed individually at maximum speed (2940 rpm) for three minutes to mix the beads, leaves and solvent and allow the beads to homogenise the plant material. One of each of the duplicate pairs of tubes containing vinegar, olive oil or butter was placed in a water bath preheated to 100°C and left to incubate for one hour; the other tube in each duplicate pair was left at room temperature for one hour. Both sets of tubes were incubated in the dark. Halfway through the incubation period, all tubes were individually briefly vortexed to mix the contents. Duplicate pairs of tubes containing only vinegar, olive oil or butter were also prepared and similarly incubated. At the end of the incubation period, all tubes were transferred to a 37°C incubator for 30 minutes to equilibrate to 37°C; this ensured that the melted butter did not solidify prior to being used for MIC testing. Nettle leaf material was removed from tubes using sterile blunt forceps, squeezing the material to release all liquid back into the tube. The liquid, containing nettle leaf extract, was used for MIC/MBC testing.

### MIC & MBC testing by broth microdilution

MIC assays using *P. aeruginosa* PA14 and *S. aureus* Newman were conducted in cation-adjusted Müller-Hinton broth (caMHB; Sigma) using Costar polystyrene tissue-culture treated 96-well plates, following EUCAST guidelines. Briefly, two-fold dilutions of each test preparation were prepared in caMHB and 50 μl aliquots of each dilution of each test preparation were added to duplicate wells containing 50 μl of each test bacterial strain suspended in caMHB at a density of ~5×10^5^ CFU/ml. Concentrations of nettle test preparations, and of vinegar, olive oil and butter, are expressed in % (vol/vol). When diluting the test preparations and adding to bacteria, vigorous pipetting was used to force an emulsion of the butter or oil to form in the growth medium. As a positive control, an MIC assay for the antibiotic meropenem (Sigma) was also conducted for both bacterial strains. Untreated control cultures were also included in the MIC plates. Plates were incubated at 37°C without shaking for 18h. Two 10 μl aliquots of each well from the MIC test plates were removed and spotted onto LB agar plus either Pseudomonas Isolation Agar (Difco; *P. aeruginosa* cultures) or Mannitol Salt Agar (supplier; *S. aureus* cultures) to select for the test bacteria and select against any endogenous bacteria present on the leaves or in the solvents. These plates were incubated at 37°C overnight (LB and PIA plates in ambient CO_2_, MSA in 5% CO_2_) and used to assess the Minimum Bactericidal Concentration (MBC) of each treatment. The remaining 80 μl of culture in each well of the MIC plate was supplemented with resazurin (supplier; filter-sterile solution in phosphate-buffered saline) to a final concentration of 45 μg.ml^−1^. The plates were shaken for 5 minutes on an orbital microplate shaker to ensure the resazurin dispersed in the wells containing oil or butter, then wrapped in foil to exclude light and incubated at room temperature for two hours. The MIC of each test substance was scored as the lowest concentration at which reduction of blue resazurin to pink resorufin was not observed. The following day, MBC plates were inspected and the MBC of each treatment was determined as the lowest concentration at which no colonies were observed. Neither the LB plates, nor the selective plates, used for MBC determination showed any visible growth of organisms other than either *P. aeruginosa* or *S. aureus*.

Pilot experiments were conducted prior to this work to assess the MICs and MBCs of several types of vinegar and olive oil to see if these varied (supplementary tables below). Red wine vinegar, white wine vinegar and cider vinegar all had low MICs/MBCs for the test species, and these were comparable between species and with MIC/MBC of 6% acetic acid, although the cider vinegar showed some variability in MIC/MBC between replicates (Table S1). We chose to work with red wine vinegar for further work. Three brands of olive oil, produced in Spain (La Española), Crete (Olive Branch) and Italy (Garofalo), all had MICs and MBC of >50%, which was the highest concentration tested (Table S2). We randomly selected Olive Branch olive oil for use in further experiments.

### Antibacterial activity of U. dioica + vinegar preparations against planktonic cultures of pathogenic bacteria

*U. dioica* leaves were immediately transferred to the laboratory and chopped roughly using a sterilised knife and chopping board. 5 g aliquots of chopped leaves were placed into 2 sterile 50 ml screw-cap tubes. 5 ml red wine vinegar (Waitrose Essentials) was added to one tube, and 5 ml sterile water to the other. For this experiment, homogenisation was carried out by hand by pounding in a sterile pestle and mortar for two minutes. Mid-log phase (6h) cultures of *S. aureus* Newman and *P. aeruginosa* PA14, grown in caMHB at 37°C, were divided into 200 μl aliquots in a 96-well microplate (Corning Costar). Aliquots of approx. 100 μl (judged by eye) of soaked leaf material was removed from each preparation of pounded nettle leaves using sterile forceps, and transferred to triplicate wells of each bacterial culture. The material was fully submerged by the culture. Further triplicate cultures were treated by adding 100 μl of either water or vinegar as negative and positive controls, respectively. The culture plates were then incubated at 37°C for 18 hours, during which time untreated bacterial cultures are expected to reach stationary phase. At the end of the incubation period, aliquots of each culture were removed, serially diluted and plated on LB agar plates. These were incubated overnight at 37°C and the number of colony-forming units (CFU) present in each culture well was calculated.

## Results

### A systematic review of published evidence for antimicrobial activity of Urtica species

A systematic review of published articles was performed in order to find primary experimental literature on the antimicrobial activity of *Urtica* extracts, as measured by the MIC or ZOI of the extract when applied to human-pathogenic bacteria or fungi, plus the model Gram-positive bacterium *Bacillus subtilis* as this is a commonly-used model organism for testing of antibacterial agents. The PRISMA diagram in Figure 1 and associated data in Datasets 1,2 and Tables S3-S5 summarise the systematic review process which led to the extraction of data from 42 publications with ZOI data and 38 publications with MIC data in our final database of eligible articles. The ZOI dataset contained many publications with missing data or low-quality data (e.g. no negative controls reported for assays; a lack of clarity about the solvent used for the final testing of extracts; unclear if assays were replicated) and a wide variety of test methods (disk diffusion and well diffusion, using different diameters of wells or disks and different thicknesses of agar; in many studies, information was lacking about well/disk diameter or agar thickness), which made it impossible to make any meaningful comparisons across studies. We thus performed no further analysis of ZOI data. Summary information and quality assessment for 38 studies containing MIC data is shown in Table 1.

**Figure 1.**
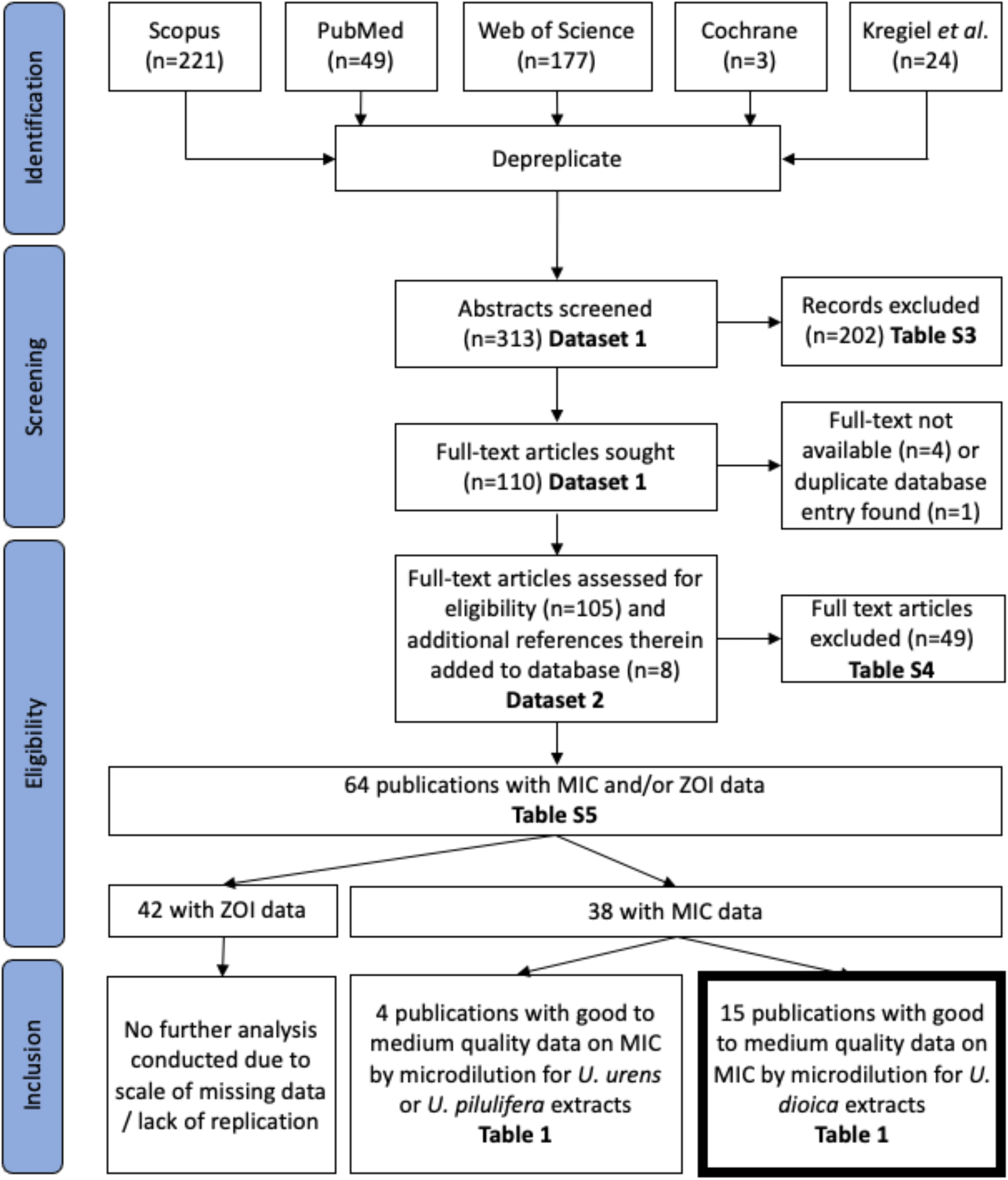
PRISMA diagram showing workflow for systematic review (19).

**Table 1.** Summary description and quality assessment of 38 publications containing eligible data on the MIC of *Urtica* spp. extracts against bacteria or fungi. Blue highlighting denotes areas where key information is missing or a quality flag, orange highlighting denotes deviations from a standard broth microdilution method. Note that some studies tested >1 strain or isolate of a species. Studies in bold were selected for inclusion in the final dataset.

| Study | *Urtica* species | Part of plant | Extract tested | Starting material | Organisms tested | MIC method | Growth medium | Temp. °C | Appropriate vehicle control? | Diagnostic guidelines referenced? | Any other quality flags | Conclusion |
| --- | --- | --- | --- | --- | --- | --- | --- | --- | --- | --- | --- | --- |
| (20) | *U. urens* | Not reported | Crude extract - ethanol | Dried | Bacteria (26 spp), fungi (16 spp) | Microdilution | Muller-Hinton broth | 37 | Yes | No | The MICs reported do not tally with the methods statement that two-fold serial dilutions of 100mg/ml stocks were used. | Exclude as concentrations reported do not make sense; also part of plant not stated |
| (21) | *U. dioica* | Not reported | Crude extract - methanol, hexane | Dried | Bacteria (6 spp), fungi (2spp) | Microdilution | Muller-Hinton / Sabouraud dextrose broth | 37 | Yes | NCCLS / CLSI | No | Exclude as part of plant not reported |
| (22) | *U. dioica* | Leaves | Crude extracts - water, ethanol | Dried |  | Microdilution | Not reported | 37 | No negative control reported, DMSO should have been used for ethanolic extract | No | No | Exclude as growth medium not reported |
| (23) | *U. dioica* | Seeds | Oils | Unclear | Bacteria (13 spp), fungi (1 sp) | Unclear | Muller-Hinton | Unclear | Yes | No | No | Exclude as MIC assay method is unclear |
| (24) | ***U. dioica*** | Whole plant | Crude extract - methanol, ethanol | Dried | Bacteria (4 spp) | Microdilution | Nutrient Broth | 37 | Yes | No | No | Include |
| (25) | ***U. dioica*** | Whole plant | Crude extract - ethanol | Dried | Bacteria (5 spp) | Microdilution | Muller-Hinton broth | 37 | No negative control reported | No | No | Include |
| (26) | *U. dioica* | Aerial parts, roots | Crude extract - water | Dried | Bacteria (4 spp) | Microdilution | Muller-Hinton broth | 37 | Yes | No | MICs of zero are reported | Exclude as reported MICs of zero do not make sense |
| (27, 28) | ***U. dioica*** | Leaves | Bioassay guided subfraction from hexane extract | Dried | Bacteria (7 spp) | Microdilution | Nutrient Broth | 37 | Used growth medium instead of DMSO as negative control | European Pharmacopeia | No | Include |
| (29) | ***U. dioica*** | Leaves | Crude extract - water | Dried | Bacteria (10 spp, fungi (10 spp) | Microdilution | Muller-Hinton / Middlebrook 7H9 broth | 37 | Used growth medium instead of DMSO as negative control | NCCLS / CLSI | No | Include |
| (30) | ***U. dioica*** | Leaves | Crude extract - ethanol | Dried | Bacteria (6 spp) | Microdilution | Muller-Hinton broth | 37 (42 for | No negative control reported | No | No | Include |

|  |  |  |  |  |  |  |  | Campylo<br>bacter) |  |  |  |  |
| --- | --- | --- | --- | --- | --- | --- | --- | --- | --- | --- | --- | --- |
| (31) | *U. dioica* | Aerial Parts | Essential oils,<br>surfactant-stabilized<br>nanoemulsions of<br>essential oils | Dried | Bacteria (3 spp),<br>fungi (1sp) | Agar dilution | Nutrient agar / Yeast<br>peptone dextrose<br>agar | 37 for<br>bacteria,<br>30 for<br>yeast | Yes | No | No | Exclude as not a microdilution<br>assay |
| (32) | *U. urens* | Leaves | Crude extract - water,<br>methanol, acetone | Dried | Bacteria (10 spp) | Agar dilution | Nutrient agar | 37 | Yes | No | It is unclear what the highest<br>concentration tested was, so we do<br>not know the meaning of "no<br>inhibition". | Exclude as not a microdilution<br>assay |
| (33) | *U. dioica* | Not reported | Crude extract - ethanol | Dried | Bacteria (5 spp) | Microdilution -<br>non-standard<br>method | Muller-Hinton broth | Not<br>reported | Unclear what<br>extracts were<br>dissolved in; water<br>used as negative<br>control | No | No | Exclude as MIC assay method was<br>non-standard; also part of plant not<br>reported |
| (34) | *U. dioica*<br>*U. pilulifera* | Seed | Oils | N/A | Bacteria (8 spp),<br>fungi (1 sp) | Microdilution | Unclear | 35 | No, only untreated<br>controls were used,<br>should have been<br>ethanol:hexane +<br>tween | NCCLS / CLSI | No | Include *U. dioica* data only |
| (35) | *U. dioica* | Aerial parts | Crude extract -<br>methanol | Dried | Bacteria (9 spp) | Well diffusion | Nutrient Agar | 37 | Yes | No | No | Exclude as not a microdilution<br>assay |
| (36) | *U. dioica*<br>*U. pilulifera* | Leaves,<br>roots, seeds | Crude extracts - water,<br>methanol | Dried | Bacteria (11 spp) | Microdilution | Muller-Hinton broth | 37 | Growth medium<br>used - ok for<br>aqueous extracts,<br>but DMSO was used<br>to resuspend<br>methanolic extracts<br>and control not<br>done | No | No | Include *U. dioica* data only |
| (37) | *U. dioica* | Leaves | Crude extract - ethanol | Dried | Bacteria (3spp) | Macrodilution | Muller-Hinton broth | 37 | Growth medium<br>used, should have<br>used methanol | No | No | Exclude as not a microdilution<br>assay |
| (38) | *U. dioica* | Leaves,<br>stems, roots | Crude extract - water,<br>ethanol | Dried | Bacteria (4 spp),<br>fungi (1 sp) | Serial dilution -<br>micro/macro<br>unspecified | Not reported | 37 | Yes | No | No | Exclude as method and medium<br>not specified |
| (39) | *U. dioica* | Not reported | Crude extracts - water, methanol | Dried | Fungi (1 sp) | Agar dilution | Sabouraud dextrose agar | Not reported | Not reported, assumed to be vehicle as this was specified for disk diffusion assay in the previous paragraph of the methods | No | No | Exclude as not a microdilution assay; also part of plant not reported |
| (40) | *U. dioica* | Aerial parts | Crude extract - hexane; Partitioned extracts - methanol, chloroform, butanol, ethyl acetate | Dried | Bacteria (6 spp), fungi (1 sp) | Macrodilution | Nutrient broth | 37 | Methanol for all, regardless of solvent | NCCLS / CLSI | No | Exclude as not a microdilution assay |
| (41) | *U. dioica* | Leaves, roots, shoots | Crude extracts - water, ethanol | Dried | Bacteria (5 spp), fungi (1 sp) | Macrodilution | Muller-Hinton broth | 37 | Growth medium used, should have been DMSO | No | No | Exclude as not a microdilution assay |
| (42) | *U. dioica* | Aerial Parts | Crude extract - ethanol, methanol | Dried | Bacteria (3 spp) | Macrodilution | Muller-Hinton broth | 37 | No negative control reported | No | No | Exclude as not a microdilution assay |
| (43) | *U. urens* | Aerial Parts | Crude extract - water, ethanol | Dried | Bacteria (8 spp) | Microdilution | Not reported | Not reported | No negative control reported | No | MIC values in the table don't match with what is said in the text; also did not report the maximum tested concentration . | Exclude due to conflicting data |
| (44) | *U. dioica* | Leaves | Crude extract - ethanol | Dried | Fungi (3 spp) | Microdilution | Modified Dixon broth | 32 | Unclear - negative control is mentioned but not specified | No | No | Include |
| (45) | *U. urens* | Leaves, stems/bark, roots | Crude extract - hexane, chloroform, ethyl acetate, methanol | Dried | Bacteria (5 spp), fungi (2 spp) | Agar dilution | Nutrient agar | 37 | Yes | No | No | Exclude as not a microdilution assay |
| (46) | *U. dioica* | Roots, stems, leaves | Partitioned extract - methanol, chloroform | Dried | Bacteria (4 spp) | Microdilution | Muller-Hinton broth | 37 | No negative control reported, should be DMSO | No | No | Include |
| (47) | *U. dioica* | Leaves | Essential oils | Dried | Bacteria (6 spp) | Microdilution | Tryptic Soy Broth | 37 | Yes | No | No | Include |
| (48) | *U. dioica* | Leaves | Crude extract - methanol | Dried | Bacteria (3 spp), fungi (1 sp) | Well diffusion | Not reported | Not reported | Yes | NCCLS / CLSI | No | Exclude as not a microdilution assay |
| (49) | *U. dioica* | Aerial parts | Crude extract - methanol | Freeze-dried | Bacteria (1 sp) | Microdilution | Muller-Hinton broth | 37 | Growth medium used, should have used methanol | No | MIC is given as mass and conversion to concentration is not clear | Exclude as MIC is not clear |
| (50) | *U. dioica* | Leaves | Bioassay guided subfraction from hexane extract | Dried | Bacteria (5 spp) | Microdilution | Nutrient Broth | 37 | Growth medium used, should have been DMSO | No | No | Include |
| (51) | *U. dioica* | Leaves | Partitioned extract - hexane | Dried | Bacteria (1 sp) | Serial dilution - micro/macro unspecified | Middlebrook 7H9 broth | 37 | Yes | No | No | Exclude as method not specified |
| (52) | *U. urens* | Whole plant | Crude extract - water, methanol | Dried | Bacteria (3 spp) | Microdilution | Muller-Hinton broth | 37 | Yes | No | No | Exclude as only two studies on *U. urens* of sufficient quality |
| (53) | *U. dioica* | Aerial Parts | Crude extract - water, methanol, ethanol, dichloromethane | Dried | Bacteria (4 spp), fungi (1 sp) | Microdilution | Not reported | Not reported | No negative control reported, also it is unclear what extracts were resuspended in | NCCLS / CLSI | No | Include, assume Muller-Hinton broth, 37°C used as diagnostic guidelines are cited? |
| (54) | *U. urens* | Leaves | Crude extract - petroleum ether; partitioned extract - ethanol; water; dichloromethane | Dried | Bacteria (6 spp), fungi (4 spp) | Microdilution | Muller Hinton / Yeast Malt / Potato Dextrose broth | 37 for Candida, 35 for others (dermatophytes) | DMSO used for all, regardless of original solvent | No | No | Exclude as only two studies on *U. urens* of sufficient quality |
| (55) | *U. dioica* | Aerial parts | Crude extract - hexane; partitioned extracts - dichloromethane, methanol, water | Dried | Bacteria (7 spp) | Microdilution | RPMI 1640 | 37 | Yes | No | No | Include |
| (56) | *U. dioica* | Leaves | Crude extract - water (comparing different methods for aqueous extraction) | Dried | Bacteria (6 spp), fungi (2 spp) | Microdilution | Muller-Hinton / Sabouraud dextrose broth | 37 for bacteria, 28 for yeast | No negative control reported, should be DMSO or methanol | No | No | Include |
| (57) | *U. dioica* | Leaves | Crude extract - ethanol | Freeze-dried | Bacteria (1 sp) | Microdilution | Muller-Hinton broth | 37 | Yes | No | No | Include |
| (58) | Not reported | Not reported | Commercial alcoholic extract | Not reported | Bacteria (1 sp) | Microdilution | Muller-Hinton broth | 37 | No negative control reported | None | No | Exclude as species and part of plant not specified |

As detailed in Table 1, there are clear gaps in the reporting of experiments and heterogeneity in the methods used. One study did not specify the method used to calculate the MIC and another used a non-standard version of the microdilution method; these two studies were both of *U. dioica* extracts and were excluded from further consideration. Of the remaining 36 studies, 26 used broth microdilution, 3 used broth macrodilution, one used an unspecified broth dilution method, two used agar well diffusion and four used agar dilution. Broth microdilution was the most widely used method, and is a standard diagnostic assay around the world. Further, an issue with agar-based methods is that some compounds cannot diffuse through the agar and so these assays are more likely than broth dilutions to give false negatives. We therefore decided to include only broth microdilution studies for further consideration. Four of the 26 microdilution studies did not specify what growth medium was used (although one of these states that NCCLS/CLSI guidelines were followed, which implies the use of Müller-Hinton broth) (16). The rest of the studies employing microdilution used a range of media, which could influence the results obtained (17). Studies were also excluded if any essential information was not reported (*Urtica* species, part of plant used, growth medium used for MIC assays, maximum concentration used in MIC assay where “no inhibition” was reported), or if the paper contained any conflicting or unclear data (examples included MIC values of zero, MIC values reported differently in data tables vs. text, and MICs given as masses not concentrations). In many cases, studies used an inappropriate negative control for MIC assays (i.e. not the vehicle in which the extract was dissolved), or did not report a negative control. However, we decided to include studies with this problem as long as they met other quality criteria, because if an extract has a MIC that is sufficiently low to be clinically meaningful, then any solvent present should be sufficiently dilute to have a negligible or no effect on bacterial growth. This process resulted in 17 studies being classified as of sufficient quality for inclusion in our review.

Of the 17 studies classified as of sufficient quality for inclusion, 15 tested extracts of *U. dioica*; as only four studies tested another species (two for *U. urens* and two for *U. pilulifera*; these last also tested *U. dioica* extracts) we focussed only on studies of *U. dioica* extracts. All of these used dried or freeze-dried plant material to make their extracts (or, in one case, seeds), but the studies used a variety of solvents and extraction techniques, plus a variety of microbial species. Twenty-six bacterial species and 11 fungal species were tested across these studies, as summarised in Table 2. To allow comparison of MIC between studies and between extraction solvents, we selected the four most commonly-studied species, which were all opportunistic bacteria that cause a range of acute and chronic infections: the Gram-negatives *E. coli*, *P. aeruginosa* and *K. pneumoniae* and the Gram-positive *S. aureus*. MIC data for these species from the included studies is contained in Table S6. One of the 15 studies (18) did not include any data on these four species, and so is not further represented in this analysis.

**Table 2.** Microbial species for which MIC assays were conducted in the 16 studies testing *U. dioica* extracts which were identified as being of sufficient quality for further analysis (see Table 1). Note that many studies tested more than one strain or isolate of the same species.

| Microbial species | n papers | References |
| --- | --- | --- |
| **Bacteria** |  |  |
| *Staphylococcus aureus* | 13 | (25),(27)/(28),(29, 30, 34, 36, 46, 47, 50, 53, 55-57) |
| *Escherichia coli* | 11 | (24, 25),(27)/(28),(30, 34, 36, 46, 47, 53, 55, 56) |
| *Pseudomonas aeruginosa* | 9 | (25),(27)/(28),(29, 34, 36, 46, 47, 50, 53) |
| *Klebsiella pneumoniae* | 8 | (24),(27)/(28),(29, 34, 36, 47, 50, 56) |
| *Bacillus subtilis* | 4 | (34, 46, 53, 56) |
| *Enterococcus faecalis* | 4 | (27)/(28),(34, 36, 47) |
| *Bacillus cereus* | 3 | (25, 30, 47) |
| *Proteus mirabilis* | 3 | (34, 55, 56) |
| *Salmonella typhi* | 3 | (27)/(28),(50, 55) |
| *Shigella flexneri* | 3 | (27)/(28),(50, 55) |
| *Acinetobacter baumannii* | 2 | (29, 34) |
| *Listeria monocytogenes* | 2 | (30, 36) |
| *Proteus vulgaris* | 2 | (36, 56) |
| *Salmonella typhimurium* | 2 | (25, 55) |
| *Streptococcus pyogenes* | 2 | (29, 36) |
| *Bacillus pumilus* | 1 | (36) |
| *Campylobacter coli* | 1 | (30) |
| *Enterococcus gallinarum* | 1 | (36) |
| *Haemophilus influenzae* | 1 | (29) |
| *Klebsiella aerogenes* | 1 | (24) |
| *Mycobacterium avium* | 1 | (29) |
| *Mycobacterium tuberculosis* | 1 | (29) |
| *Salmonella infantis* | 1 | (30) |
| *Serratia marcescens* | 1 | (24) |
| *Shigella sp.* | 1 | (36) |
| *Staphylococcus epidermidis* | 1 | (29) |
| *Streptococcus pneumoniae* | 1 | (29) |
| **Fungi** |  |  |
| *Candida albicans* | 4 | (29, 34, 53, 56) |
| *Candida parapsilosis* | 2 | (29, 34) |
| *Aspergillus niger* | 1 | (56) |
| *Candida krusei* | 1 | (29) |
| *Candida tropicalis* | 1 | (29) |
| *Epidermophyton floccosum* | 1 | (29) |
| *Malassezia furfur* | 1 | (44) |
| *Malassezia globosa* | 1 | (44) |
| *Malassezia slooffiae* | 1 | (44) |
| *Microsporum gypseum* | 1 | (29) |
| *Trichophyton rubrum* | 1 | (29) |

The MIC values obtained from the 14 selected studies for the four chosen bacterial species are shown in Figure 2, separated by the type of solvent/extract tested and colour coded by the part of the plant used to make the extract. Figures S1 and S2 show the same data colour coded by the strain of each species used, and by the study from which the data was taken, respectively. While some studies in our systematic review reported a promisingly low MIC of some *Urtica* extracts, most did not. MICs were often > 100 μg.ml^−1^ and there was a lot of variation in MICs obtained for plant material extracted in the same solvent by different authors. Further, the studies varied widely in quality and it is very hard to obtain a clear picture of the potential for antibacterial activity as such a variety of plant parts, solvents and bacterial strains were used. Overall, despite some promisingly low MICs (occasional aqueous or ethanolic extracts, essential oils, seed oils and hexane subfraction of leaves) there is not a strong suggestion of antibacterial potential in this dataset.

**Figure 2.**
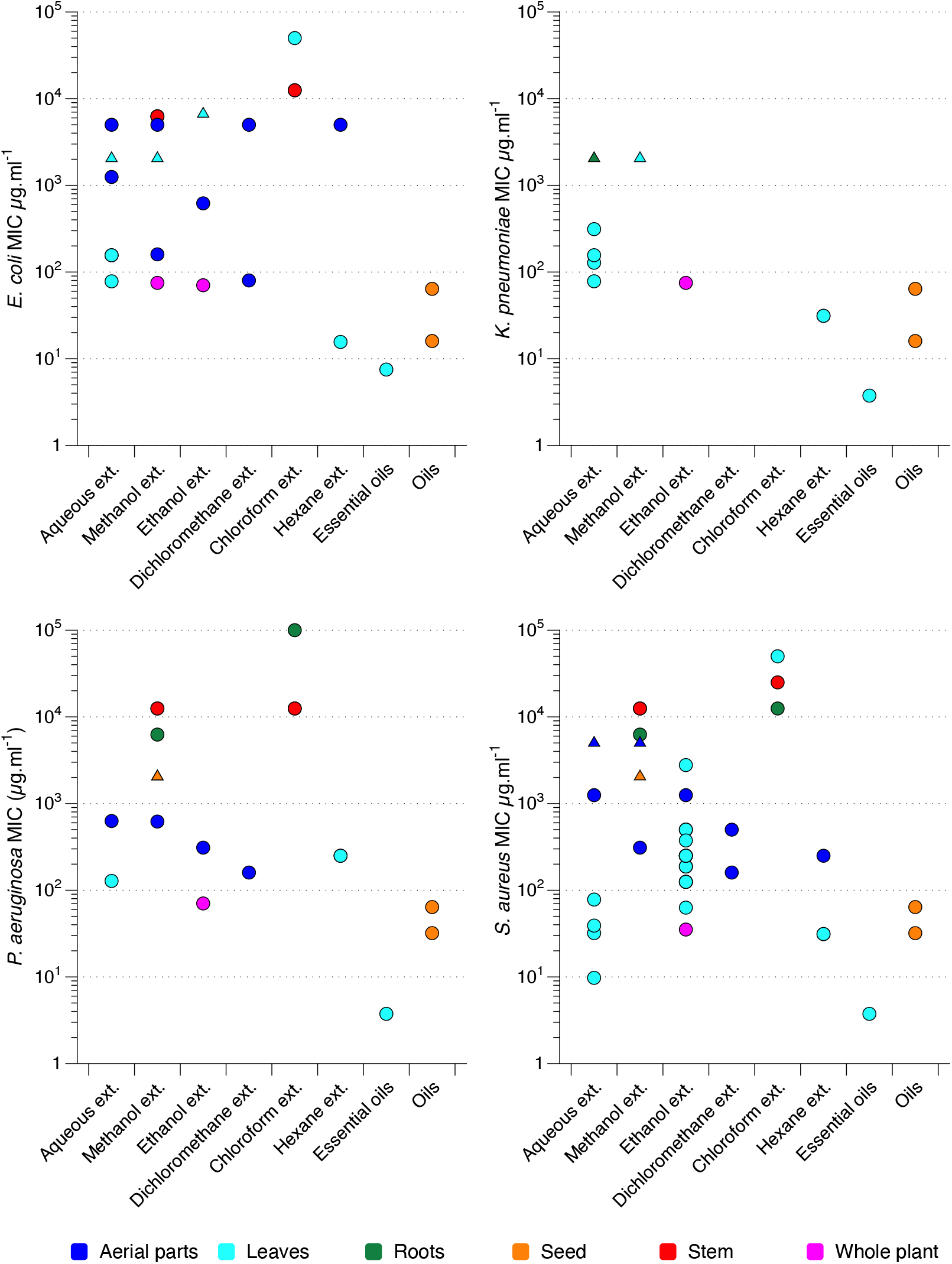
MIC values for nettle extracts tested against various strains and isolates of *E. coli*, *P. aeruginosa*, *K. pneumoniae* and *S. aureus*, extracted from studies described in Table 1; raw data are provided in Table S4. Each circle denotes the MIC value for an individual bacterial strain or isolate, obtained from one study. Triangles denote where the MIC is greater than the highest concentration tested, the MIC is therefore presented as the highest concentration tested. The *x* axis records the type of solvent (water, methanol, ethanol, dichloromethane, chloroform, hexane) used to make the extract (ext.), or the class of compounds extracted (essential oils, oils). Note that exact extraction methods varied between studies. The *x* axis is ordered with more hydrophilic/lipophobic extracts on the left, and more hydrophobic/lipophilic extracts on the right. The same data are presented again with different colour coding in Figure S1 to show the different strains or isolates of each bacterial species used, and in Figure S2 to show which data points came from which study.

The three clinical trials of nettle extracts to treat infection which were found in the Cochrane Central Register of Controlled Trials, and which did not provide any MIC data, also did not report strong evidence for a beneficial effect of the extracts used (59–61).

It was striking on reading the articles used in this review that the authors of experimental studies never explicitly linked the methods they used to historical or traditional methods of preparing nettles for medicinal use, for example, by talking about how the material is traditionally prepared or applied/taken by the patient. Almost all introductions or discussions connected the reason for the research with a long tradition of use (“since the dawn of time” terminology) or popularity in specific folk medicines. The reports often pointed to the conditions being treated, but none discussed how the plant is prepared in those traditional or historical methods, or if those preparation methods informed the authors’ choices of plant parts and extraction methods. This is important because a preparation where the plant is traditionally boiled in water and drunk will extract different natural products from the plant than a preparation where the plant is pounded in fat or oil to produce an ointment, and tests of each use should ideally recapitulate the traditional preparation process in the lab (i.e. in this case using hydrophilic vs. lipophilic solvents).

Of direct relevance to the pre-modern European wound preparations outlined in the Introduction, we noted a relative dearth of studies looking at lipophilic extracts, and we did not see any studies which paralleled the use of vinegar (aqueous extract but in a low pH solvent). The two studies which did look at lipophilic extracts did report some of the lowest MICs in our dataset. These were seed oils (34) and essential oils (62); note “essential oils” are volatile oils and the study which extracted these from nettle leaves reported a preponderance of fatty acids in the extract. The lack of focus on fats/oils is even more interesting because of reports that lipophilic extracts of nettle have good anti-inflammatory activity (63). The potential use of vinegar is especially intriguing because vinegar itself is a potent antimicrobial – acetic acid at concentrations below that present in vinegar can kill bacteria growing as biofilms, and is currently the subject of a clinical trial for the treatment of antibiotic-resistant burn wound infections (11–13). Thus we must consider the potential that nettle-derived natural products with weak antibacterial activity on their own may synergise with an antibacterial agent like vinegar to produce a mixture with enhanced activity (64, 65). To address this gap in the published data, we conducted MIC testing of fresh *U. dioica* leaf extracts prepared in vinegar, olive oil or butter.

### Extracts of U. dioica leaf prepared in vinegar, butter or olive oil do not inhibit the growth of P. aeruginosa or S. aureus in a broth microdilution assay

We made extractions of fresh *U. dioica* leaves in red wine vinegar, butter or olive oil and tested the extracts for their MIC against *P. aeruginosa* PA14 and *S. aureus* Newman in a microdilution assay in cation-adjusted Müller-Hinton broth. We compared extracts made at room temperature with extracts made by boiling the leaves + solvent mixtures. Vinegar, olive oil and butter were also assessed for their MIC as negative controls. We followed EUCAST guidelines for MIC testing (66) with the exception that we added resazurin to cultures at the end of the assay to assess whether cultures contained live bacteria or not; this was because of the presence of sediment from the natural products used, and the colour of the preparations, which made scoring turbidity of cultures by eye difficult. Minimum bactericidal concentrations were also assessed by plating out an aliquot of the MIC plate prior to the addition of resazurin.

As shown in Table 3, there was no inhibitory effect of the nettle leaf extracts at all, regardless of solvent and regardless of whether the extraction was conducted at 100°C or room temperature. On the whole, the nettle leaf extract did not interfere with the antibacterial activity of vinegar. The exception to this was the case of the 100°C preparations tested against *P. aeruginosa*, where the presence of nettle extract increased the MBC of vinegar from 3.125% to 12.5%. (A >2-fold change in the MIC or MBC is generally considered biologically meaningful).

**Table 3.** MICs and MBCs of *U. dioica* leaf extracts against *P. aeruginosa* and *S. aureus*. Where a single value is reported, both duplicate assays returned this value. Where two values are reported, the duplicate assays gave different results.

|  | Test agent | S. aureus Newman |  | P. aeruginosa PA14 |  | Units |
| --- | --- | --- | --- | --- | --- | --- |
|  |  | MIC | MBC | MIC | MBC |  |
|  | Meropenem | 1 | 1 | 0.125 | 2 | µg.ml<sup>-1</sup> |
| Extraction for<br>1 hr at room<br>temperature | Vinegar | 3.125 | 6.25 | 1.56 | 3.125 | % vol/vol |
|  | *U. dioica* vinegar extract | 3.125 | 6.25 | 3.125 | 6.25 |  |
|  | Olive oil | >50 | >50/50 | >50/25 | >50/50 |  |
|  | *U. dioica* olive oil extract | >50 | >50 | >50 | >50 |  |
|  | Butter | 25 | >50 | >50/50 | >50/50 |  |
|  | *U. dioica* butter extract | 50 | >50 | >50 | >50 |  |
| Extraction for<br>1 hr at 100°C | Vinegar | 3.125 | 6.25/12.5 | 1.56 | 3.125 |  |
|  | *U. dioica* vinegar extract | 6.25 | 12.5 | 3.125 | 12.5 |  |
|  | Olive oil | 50 | >50/50 | >50 | >50/50 |  |
|  | *U. dioica* olive oil extract | 50 | >50 | >50 | >50 |  |
|  | Butter | >50/50 | >50 | >50 | >50 |  |
|  | *U. dioica* butter extract | 12.5 | >50 | >50 | >50 |  |

### *U. dioica* leaves act as a sponge to transfer bactericidal concentrations of vinegar to planktonic cultures of *P. aeruginosa* and *S. aureus*

While performing this work, we noticed that the nettle leaves were highly absorbent – 2.5 g of chopped leaves readily absorbed vinegar when pounded, and the leaves had to be squeezed very firmly to release the liquid. We therefore hypothesised that the use of nettles in historical remedies may be as a relatively safe natural substance which could be employed as a sponge to deliver useful agents such as vinegar to a wound or other infection site. To test this hypothesis, we treated triplicate mid-log phase cultures of *P. aeruginosa* and *S. aureus* with 0.5 vols of nettle leaves that had been pounded in 0.5 vols of sterile water alone or 0.5 vols of red wine vinegar alone. To better approximate historical methods of remedy preparation, instead of using homogenisation beads we pounded the nettle leaves in water or vinegar by hand using a mortar and pestle. This confirmed immediately that the nettle leaves were highly absorbent: water or vinegar very quickly absorbed into the leaves, leaving no visible liquid in the mortar. Only sub-millilitre amounts of liquid could be released from the mass of leaves on pounding with the pestle once the liquid had been absorbed. As shown in Figure 3, the nettles had no effect on viable bacteria numbers themselves, but were able to carry sufficient vinegar with them to completely eradicate the bacterial populations (all replicates of nettle + vinegar and vinegar treatments for both species had viable cell numbers below the limit of detection by plating).

**Figure 3.**
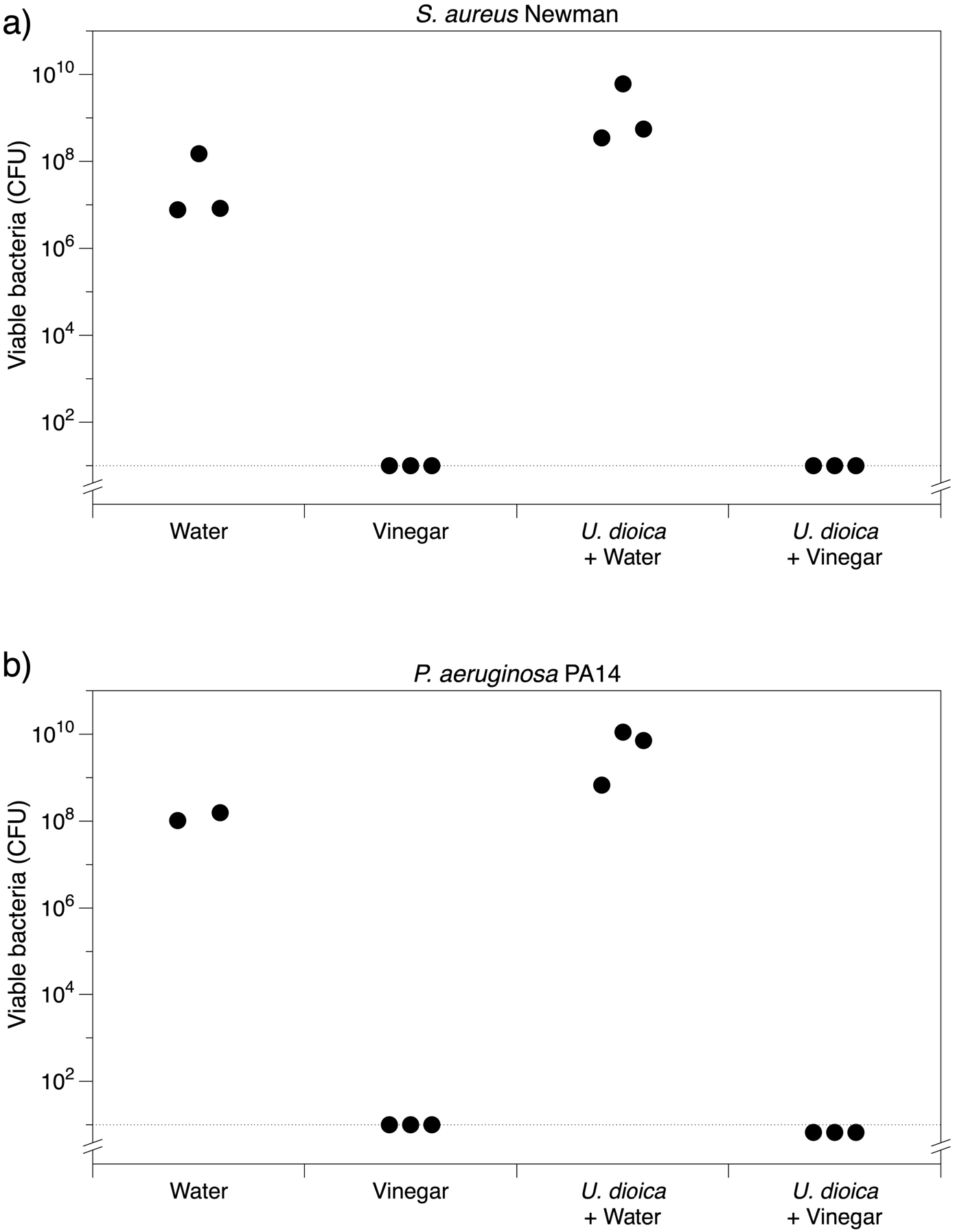
The effect of U. dioica leaves, chopped and pounded in either water or vinegar, on viable bacterial counts (colony-forming units, CFU) in cultures of (a) *S. aureus* Newman and (b) *P. aeruginosa* PA14 after 18h exposure.

## Discussion

The systematic review revealed hundreds of published studies on the antimicrobial properties of *Urtica* species. However, the quality and techniques used are too variable to draw meaningful conclusions. In a selected set of decent quality studies looking at MIC of *U. dioica* extracts against bacteria, we found a promisingly low MIC of some extracts, but most MICs were >> 100 μg.ml^−1^. There is a good representation of extraction techniques employing ethanol or water, but there is almost no representation of oils and fats, which were commonly used in the preparation of nettles in our team’s study of select medieval medical preparations. Due to time constraints, only one aspect of the potential bioactivity of nettles (antimicrobial) could be considered for the systematic review. In future, it may be worth performing a similar review of experimental data on the anti-inflammatory or immuno-enhancing capabilities to follow up reports of effective anti-inflammatory activity of lipophilic extracts of nettle (63, 67).

The experimental results from our study of nettles+vinegar, nettles+oil and nettles+butter preparations showed that these treatments did not extract anything antibacterial from the nettles. However, the nettles acted as a highly-effective sponge and delivery agent for vinegar, which is very well established as a bactericidal agent (11–13). Many of the historical remedies that we identified in our dataset of historical medicines are poultices (8). It is possible that nettles were used in the past at least partly for their ability to enable large volumes of medicinal liquid to be delivered topically to a wound or ulcer and remain securely in place. The use of fresh nettles leaves as a sponge is a stark contrast with modern attempts to extract molecules from dried leaves: all the papers in our systematic review used dry plant materials. The sponge-like qualities and ability to carry vinegar, for instance, may make nettles an excellent wound dressing.

A limited number of studies on the anti-inflammatory or immunomodulatory effects of nettle extracts have suggested that this was due to the sting triggering an antihistamine response (68). Broer & Behnke reported that an isopropanol extract from nettle leaves reduced primary T-cell responses, which could have an immunomodulatory effect in T-cell mediated diseases, such as rheumatoid arthritis (69). Isopropanol is a relatively hydrophilic solvent. In contrast, Johnson et al. found that their dichloromethane extract, which is lipophilic, had the best combination of low cytotoxicity and good anti-inflammatory activity (as measured by its ability to reduce LPS-mediated activation of NF-κβ) (63). It is important to note that these results do not mean that historical or traditional recipes produced the same effects.

The body of literature available suggests that lipophilic extracts of nettles seem to be promising for the purposes of extracting molecules which could be used in purified and concentrated formulations for drug development; while our results indicate that the function of nettle plant material in traditional topical preparations for wounds may simply be as a poultice medium for holding antibacterial or soothing ingredients at the treatment site. In our team’s dataset of medieval remedies, nettles are frequently combined with fats and oils. Fats and oils probably functioned primarily as vehicles to convey medicines, as a palliative agent due to their emollient properties, or to absorb blood or restrain bleeding in a wound. However, the possibility that lipophilic extracts of nettle may have anti-inflammatory activity, and that preparing nettles in culinary fats and oils might produce anti-inflammatory extracts, would be an interesting area for future research.

It may also be worth noting that, as in the recipe examples given in the introduction, a number of known antimicrobial medicinal plants and products are combined with the nettles in our dataset of medieval remedies. It is possible that additional biological activity is achieved when *Urtica* spp. are combined with other natural products, due to additive or synergistic interactions between molecules from different plants (70, 71).

Perhaps the overall message from this work is that sometimes, when studying historical/traditional uses of plants as medicines, we should “keep it simple”. While ethnopharmacological research has discovered that some plants used in historical treatments have clinically significant bioactivity, others may have been employed simply as useful physical components of a remedy to facilitate the delivery of other ingredients to the treatment site.

---

**Figure S1.**
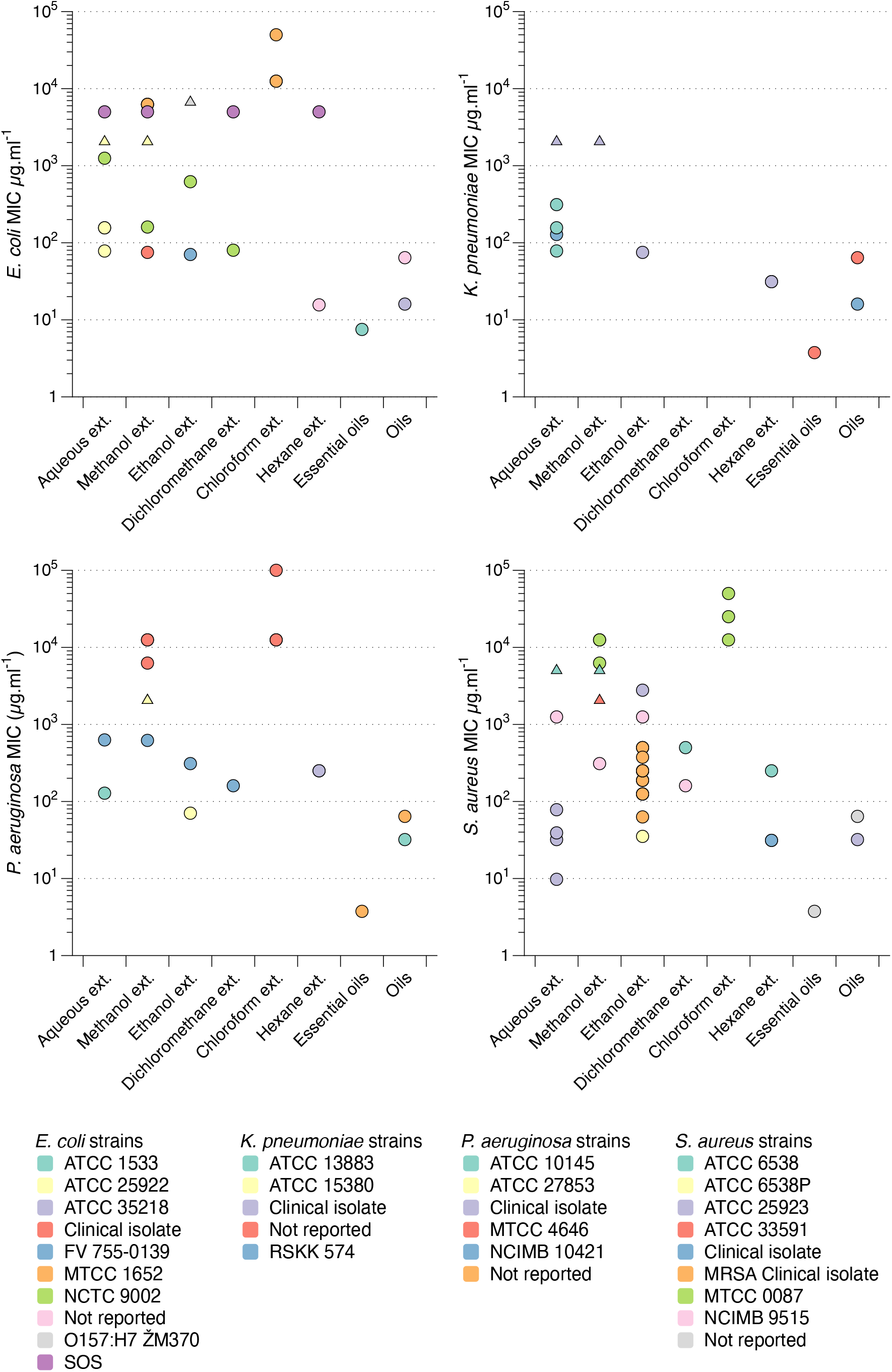
Data from Figure 1, colour coded by bacterial strain or isolate tested.

**Figure S2.**
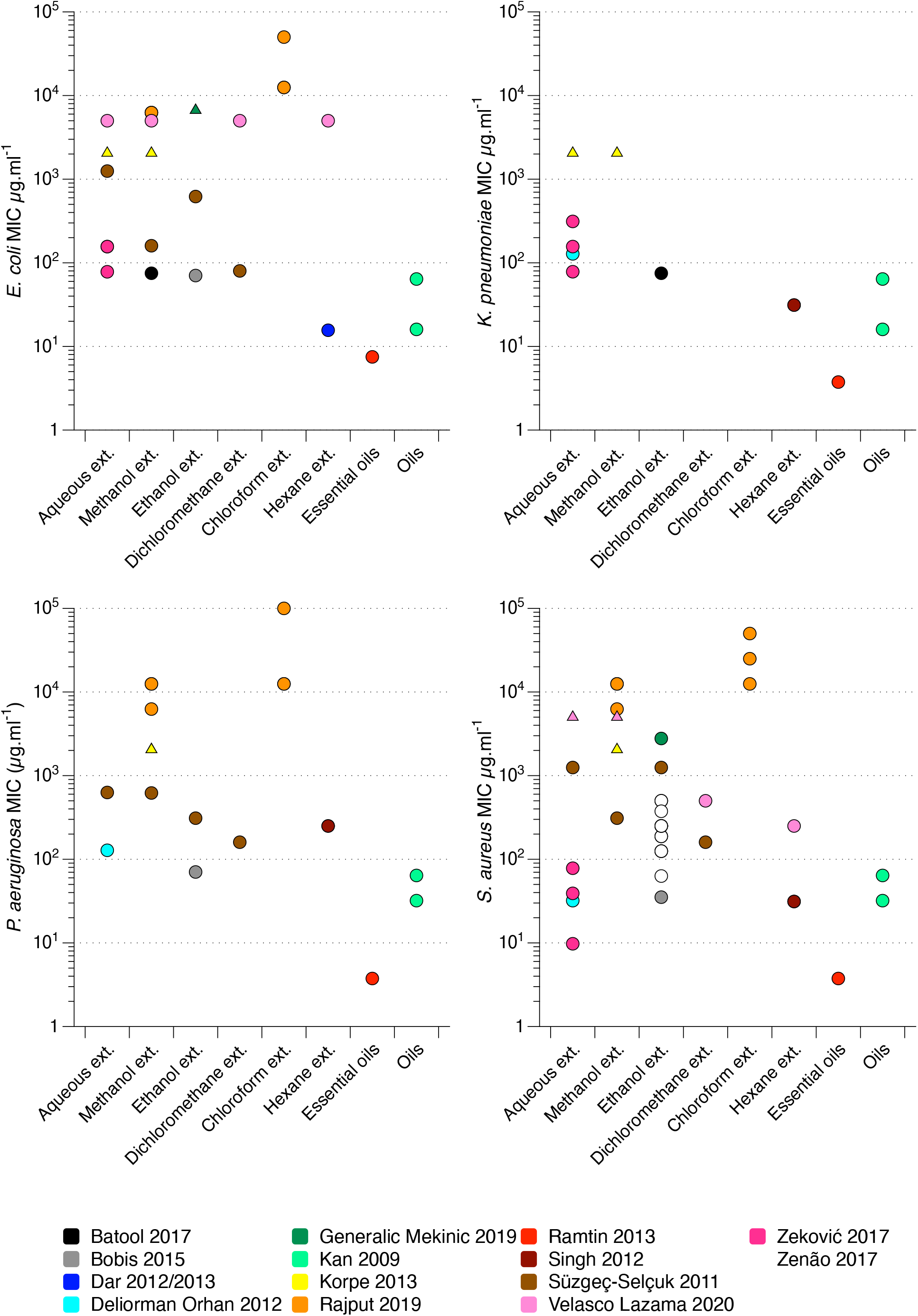
Data from Figure 1, colour coded by the study from which the data point was extracted.

## Author statements

Conceptualisation: FH, EC

Investigation: FH, EC, JFP

Data curation: FH, EC

Writing – Original Draft Preparation: FH, EC

Writing – Review and Editing: FH, EC, JFP

Funding: FH, EC

## Acknowledgements

We would like to thank Dr Christina Lee for orchestrating and leading the overall “Nettles and Networks” project. The team would like to thank Dr Sheyda Azimi, who kindly read and extracted data from two articles written in Farsi; Prof. Orkun Soyer, who kindly read two articles written in Turkish; and Kate Willett, who helped conduct lab work during her University of Warwick URSS summer vacation project. We also extend thanks to Dr Diana Luft for sharing her translations of medieval Welsh recipes. This study was aided by a pilot project on nettles and vinegar undertaken by Navneet Jandu and funded by the Microbiology Society’s Harry Smith Vacation Studentship. We were also aided by Conor Macnab’s final-year BSc project in the School of Life Sciences at the University of Warwick, which showed there was sufficient published data on nettle extracts to carry out the larger-scale efforts of our team’s project.

